# Exposure to artificial light at night accelerates but does not override latitude-dependent seasonal reproductive response in dark-eyed juncos

**DOI:** 10.1101/2020.06.21.163774

**Authors:** D. Singh, J. Montoure, E.D. Ketterson

## Abstract

In the modern era of industrialization, illuminated nights have become a common defining feature of human-occupied environments, particularly cities. Artificial light at night (ALAN) imposes several known negative impacts on the neuroendocrine system, metabolism, and seasonal reproduction of species living in the wild. However, we know little about the impact of ALAN on populations of birds that either live year-round in the same location or move to different latitudes across seasons. To test whether ALAN has differing impact on reproductive timing of the bird populations that winter in sympatry but breed at different latitudes, we monitored sedentary and migratory male dark-eyed juncos that were or were not exposed to low intensity (~2.5 ± 0.5 lux) ALAN. All groups were held in common conditions and day length was gradually increased to mimic natural day length changes (NDL). We assessed seasonal reproductive response from initiation to termination of the breeding cycle. As expected based on earlier research, the sedentary birds exhibited earlier gonadal recrudescence and terminated breeding later than the migratory birds. In addition, resident and migrant birds exposed to ALAN initiated gonadal recrudescence earlier and terminated reproduction sooner as compared to their conspecifics experiencing NDL. Importantly, the difference in the reproductive timing of sedentary and migratory populations was maintained even when exposed to ALAN. This variation in the seasonal reproductive timing may likely have a genetic ground or early developmental effects imposed due to different latitude of origin. This study reveals first that latitude-dependent variation in reproductive timing is maintained despite exposure to ALAN, and second that ALAN accelerated reproductive development across both migrants and residents. The results corroborating relationship between latitude, population, and ALAN impact on seasonal reproductive timing, may provide a potential mechanism to test the fitness of a population and its range expansion to exploit urban environment.

## Introduction

Urbanization is transforming landscapes across the globe in part as a severe consequence of anthropogenic light pollution (Shochat et al., 2006; Falchi et al., 2016; Kyba et al., 2017). Sources of artificial light at night (ALAN) are a potential threat to biodiversity by interfering with a variety of seasonal behaviors such as migration, hibernation, and reproduction (Longcore and Rich, 2004; Miller, 2006; Rodriguez and Rodriguez, 2009; Kempenaers et al., 2010). Natural variation in photoperiod (day length) is known to provide a consistent and accurate cue to coordinate seasonal events defined as distinct life-history states (LHSs) in an annual window. Reproduction in birds is strictly regulated by photoperiod in the majority of species breeding at mid to high latitude (Wingfield et al., 1992; Bronson and Heideman, 1994; Dawson et al., 2001). Photoperiod serves as the primary predictive cue triggering annual gonadal recrudescence and regression in seasonally breeding animals, which also rely on supplementary cues to initiate and regulate timing of reproductive development (Bronson and Heideman, 1994; Dawson 2001; Wingfield 2012). When gonads recrudesce in spring, the rate of gonadal growth has been shown to be directly proportional to increasing photoperiod (Farner and Wilson, 1957; Follett and Maung, 1978). Later towards the end of the breeding season, while days are still long, many bird species reduce gonad size and circulating testosterone and become photorefractory (Burger, 1949; Miller, 1954). Exposure to short day lengths during autumn breaks photorefractoriness and restores a bird’s ability to develop gonads in response to increasing day lengths the following spring (Farner and Mewaldt, 1955).

Different species match their life-history states to the availability of food supplies at different times of the year, hence the optimal time for breeding varies by species and even populations within species (Dawson and Goldsmith 1983; Wingfield et al., 1992; Dawson et al., 2001; Ball and Ketterson 2008; Watts et al., 2015). For example, the duration of different life history states varies in birds breeding at different latitudes to match their breeding and nestling growth to the peak resource availability time (Lack 1968; Visser at al., 2004). Any mismatch between timing of breeding, nestling growth, and availability of required food resources could have deleterious effects on the population (Visser at al., 2004; Jonzén et al., 2006). In general, birds living at lower latitudes initiate gonadal growth earlier in the spring when days are shorter, remain in the stimulatory phase for longer, and terminate breeding later, as compared to birds that breed at higher latitudes (Dawson 2001; Singh et al., 2019).

While many studies have addressed population differences in timing of initiation and termination of reproduction in populations that breed at different latitudes in natural day light cycle, a significant knowledge gap remains as to whether and how timing differences among populations are affected by ALAN. Just as seasonal responses to natural day lengths are driven by the hypothalamic-pituitary-gonadal (HPG) axis (Partecke et al., 2005; Dominoni et al., 2013a), the same may be true for ALAN. Gonadotropin releasing hormone 1 (GnRH1) released from the hypothalamus is a key regulator that stimulates the release of the gonadotropins, luteinizing hormone (LH) and follicle-stimulating hormone (FSH), from the pituitary (Li et al., 1994; Cho et al., 1998). LH and FSH communicate to target tissues, the gonads, and start development of gametes along with release of sex steroids.

The impacts of ALAN on seasonal reproduction have been studied in bird species under controlled laboratory conditions as well as in the field (Bruening et al., 2016; Dominoni, 2015; Gaston et al., 2017; de Jong et al., 2015; Robert et al., 2015). In a lab study, European blackbirds (*Turdus merula*) exposed to ALAN (0.3 lux) showed 3 weeks advanced gonadal growth as compared to their conspecifics exposed to dark nights (Dominoni et al., 2013a). A field study on great tits (*Parus major*) areas with ALAN advanced egg-laying date (de Jong et al., 2015) as compared to birds experiencing dark nights. In addition, female blue tits (Cyanistes careuleus) breeding in close proximity of street lights laid eggs 1.5 days in advance of females breeding in dark areas (Kempenaers et al., 2010). Other field studies have shown that ALAN can extend foraging activity (Russ et al., 2015), advance dawn singing times (Kempenaers et al., 2010), and increase daily locomotor activity, with early onset and delayed offset activity time associated to ALAN exposure (Dominoni et al., 2013; de Jong et al., 2016). Given that the majority of animals across the globe use of photoperiod to time reproductive development in anticipation to the expected maxima for annual resource availability, the effects of ALAN exposure on reproduction are likely to be far-reaching (Bradshaw & Holzapfel, 2010; Helm et al., 2013).

This study reports the impact of ALAN on a songbird, the Dark-eyed Junco (*Junco hyemalis*). The species consists of a complex of sub-species and populations; some are migratory and some are sedentary (i.e., resident) During winter and early spring, migrants and residents are found in sympatry (Fudickar et al., 2016; Grieves et al., 2016). As day length increases, residents initiate gonadal growth prior to the departure of migrants to the North, indicating a differential response to the same photoperiod in spring. In a recent study, when held captive in a common garden and exposed to gradually increasing photoperiod simulating the natural increase in photoperiod resident and migrant male juncos exhibited different critical photoperiodic thresholds (CPP) (Singh et al., 2019). Residents, which breed at lower latitudes, expressed indices of reproductive preparedness earlier than migrants, i.e. they exhibited a lower CPP (Singh et al., 2019).

Here we asked how ALAN might affect the reproductive endocrine response in songbird populations that co-exist but differ in the timing of reproduction. We employed external GnRH challenge a tool that can be used to investigate precise variation in animals behavior and physiological state. We administered controlled doses of GnRH (i.e., a GnRH challenge) to individuals and measured elevated T in response to external GnRH challenge (T_GnRH_) as a downstream response by the gonad (Jawor et al., 2006; Spinney et al., 2006; Grieves et al., 2016) to obtain a measure of the gonadal response (Singh et al., 2019). We exposed groups of resident and migrant juncos to either normal photoperiods or ALAN under controlled laboratory conditions. Given that ALAN can be perceived as extended photoperiod by the neuroendocrine system, we predicted that exposure to ALAN would advance the growth of the gonads, elevate T_GnRH_, and lead to earlier initiation of reproduction than those exposed to a normal photoperiod. More specifically, we predicted that ALAN would accelerate the gonad growth rate i.e., stimulate earlier rise in T_GnRH_ and lead more rapidly to peak stimulation as compared to conspecifics exposed to natural day length (NDL). We did not have an a priori expectation about how ALAN would influence the earlier observed difference in photoperiodic threshold between migrants and residents. One outcome might be that ALAN would accelerate recrudescence in both groups but the gap between them would be maintained. That is, both groups would accelerate gonadal growth but residents would still respond earlier than migrants. If the migrant and resident birds exposed to ALAN were to maintain the population-level differences in their reproductive response, this would provide strong support for genetic (or early developmental) determination of how the HPG axis responds to light. If instead exposure to ALAN were to eliminate the timing difference between migrants and residents, the implication would be phenotypic plasticity that overrides population-level differences in seasonal timing.

Based on earlier comparisons of migrants and residents, we also anticipated that birds exposed to ALAN would terminate gonad growth sooner and undergo molt earlier than birds exposed to NDL. Prior studies with GnRH challenges have been based on challenges conducted during the day, but many hormones including testosterone show daily variation mediated by glucocorticoids and mineralocorticoids (Harden et al., 2016). To explore daily variation in circulating T levels, we measured day and night testosterone. Considering the ALAN impact on daily clock we predicted that day and night T_GnRH_ levels would differ in comparison to birds exposed to dark night. ALAN imposes negative impact on the reproductive axis by elevating corticosterone (CORT) levels (Wingfield and Sapolsky 2003). However, a study on dark eyed junco found substantially higher level of night time testosterone (Needham et ail., 2017). In this paper, we provide results showing relationship between increased ALAN impact on reproductive timing of birds that breed at different latitude and have different endocrine response.

## 1. Methods

### (a) Study Species

The dark-eyed junco (*Junco hyemalis*) is a passerine distributed broadly across North America (Nolan et al., 2002) with populations that diverged approximately 15,000 years ago into multiple subspecies that differ in migratory behavior, reproductive timing, plumage coloration, and, morphology (Milà et al. 2007; Atwell et al., 2011; Fudickar et al., 2016). Within this species complex, a migratory subspecies, *J. h. hyemalis,* (hereafter ‘migrants’) breeds in temperate coniferous and mixed forests across Canada and Alaska, whereas a sedentary subspecies, *J. h. carolinensis*, (hereafter, ‘residents’) is found year-round in the Appalachian Mountains of the eastern United States. The migratory and resident subpopulations are found in overlapping distributions during the winter from October through late March. In spring migrants start departing the wintering ground at the same time that residents begin to grow gonads and form pair bonds (Fig 1a).

**Figure 1.**
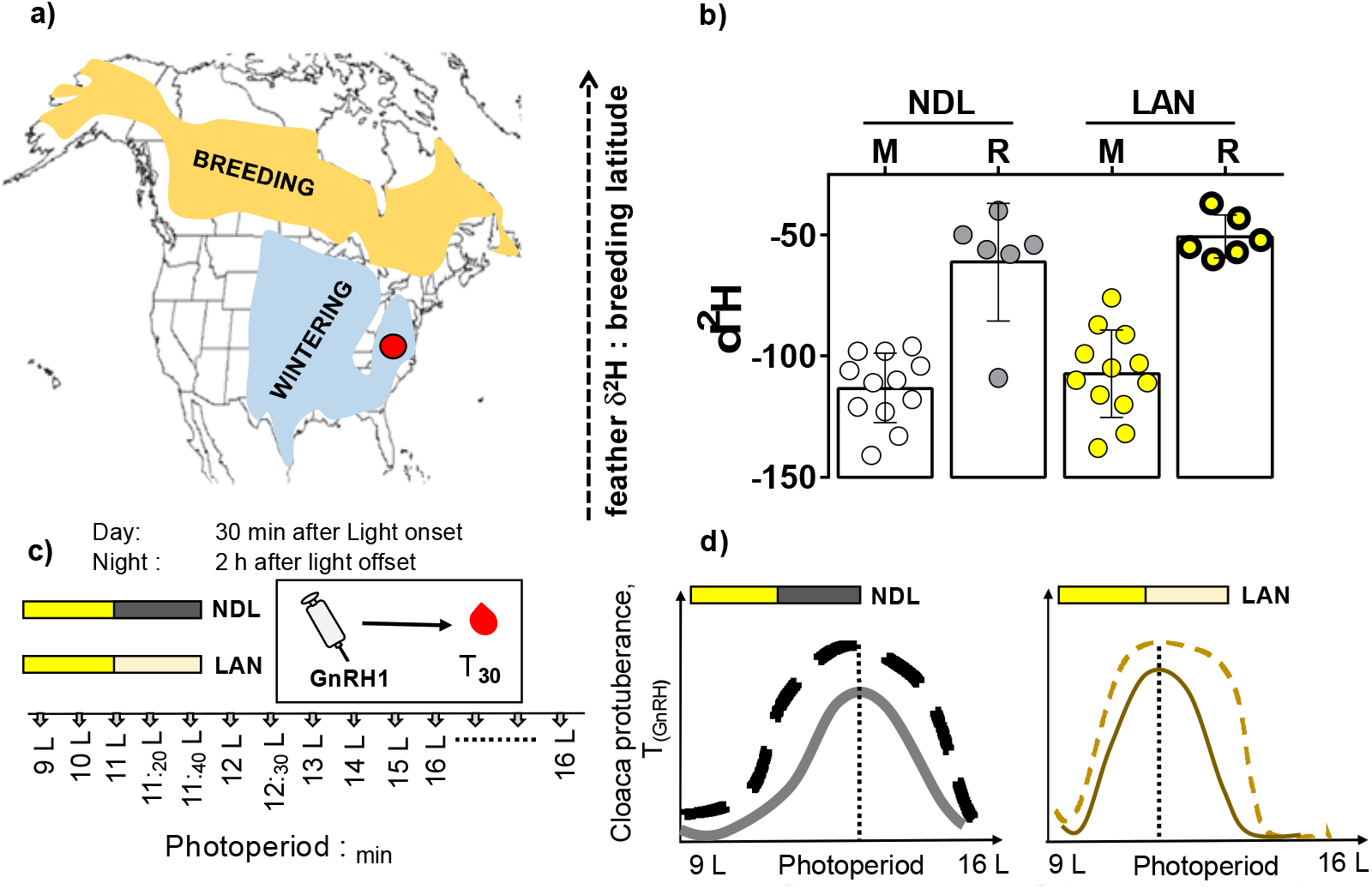
Schematic showing geographical distribution of resident (*J. h. carolinesis*; red circle; breeds year round on the same ground) and migrant (*J. h. hyemalis*; orange as breeding area, pale blue as wintering area) on the North America map (a). The upper right panel represents the feather δ^2^H distribution of resident and migrant juncos under different light treatments. The bar graph represents mean difference in resident and migrant juncos’ hydrogen isotopes. Values of δ^2^H on the Y-axis are represented as hollow circle for migrants under natural day length (LAN), solid grey circle for residents exposed to natural day length (NDL), solid yellow circle for migrants exposed to light at night (LAN), and thick solid yellow circle for residents exposed to LAN (b). Lower left panel showing the experimental protocol in which resident and migrants were divided in two light treatments, one had increasing photoperiod with dark nights and other group had similar photoperiod except nights were illuminated with dim light (c). All physiological parameters were measured along with GnRH challenged blood to measure T levels in increasing photoperiod. Prediction for reproductive response in resident and migrant juncos exposed to different light treatments (d).

### (b) Bird Capture and Housing

Adult male sympatric resident (n=18) and migrants (n=22) male dark-eyed juncos were captured in the month of December, 2017 at University of Virginia’s Mountain Lake Biological Station in Giles County (37.37 °N, 80.52°W). The resident and migrant birds were differentiated based on bill coloration (pink bill, migrant; gray bill, resident: Nolan 1976), body size, and difference in the body mass (resident birds are bigger and heavier than migrants) during non-reproductive state (Pyle, 1997; Fudickar et al., 2016; Singh et al., 2019).

All the birds were transported to Kent Farm Research Station (KFRS) in Bloomington, Indiana and housed in the indoor aviary with *ad libitum* food and water. These birds experienced gradually increasing then declining photoperiod simulating annual day length cycle to determine the photoperiodic threshold for gonad recrudesce response from the beginning till the end of year 2018 (Singh et al., 2019). All birds had already completed one reproductive cycle in captive condition and were maintained in photosensitive state at 9h day length till the start of this experiment testing impact of artificial light at night on the reproductive timing in the year 2019. All collection and sampling appropriate scientific collecting permits issued by the Virginia Department of Game and Inland Fisheries (permit # 052971) and the US Fish and Wildlife Service (permit # 20261).

### (c) Feather Stable Hydrogen Isotopes

We used both morphology and δ^2^H composition of all birds’ feathers to classify the resident from migrant population (Fig 1 b; Fudickar et al., 2016; Singh et al., 2019). We used the δ^2^H values for respective individuals (reported in Singh et al., 2019) as a continuous variable against all the response variable.

### (d) Light treatments

In order to monitor differences in gonadal recrudescence and related physiological changes between residents and migrants in response to increasing day length and constant dim artificial light at night (~2.5 ± 0.5 lux), we artificially regulated changes in day length while exposing one treatment group to total darkness and night and one to constant ~2.5 ± 0.5 lux average intensity throughout the night. To determine intensity in each compartment, we averaged light intensity from all the four corners of the room and along different levels top, middle and bottom of the trees birds used to perch in during the average day (140 ± 5 lux) and night (~2.5 ± 0.5 lux). We used LI-COR light meter with LI-190R quantum sensor that measures visible range photosynthetically active radiation per unit area (PAR, in μmol of photons m^2^/s). The PAR value was converted to the lux intensity for white light emitting diode (LED) light source using a horticulture online calculator (https://www.waveformlighting.com/horticulture/convert-ppfd-to-lux-online-calculator). In addition, birds were exposed to, very dim red light source headlamps when they were sampled at night (see below). Photoperiod was increased every fourteen days from February 16 to September 2 in the following schedule: 9L:15D (starting point when all birds were experiencing 9h of day light and dark night), 9L:15D (1; birds were separated in two separate groups one with dark night and other with constant dim artificial light at night), 10L:14D, 11L:13D, 11:_20_L:12._40_D (11h and 20 minutes of light and 12h and 40 minutes of dark), 11:_40_L:12:_20_D, 12L:12D, 12:_30_L:11:_30_D, 13L:11D, 15L:9D, 16L:8D (Fig 1c).

We then continued all observations for 8 weeks in 16L:8D. Hereafter, each photoperiod schedule is referred to as the number of hours: minutes of light. At the end of each photoperiod, birds were bled and processed for physiological measurements. To study the daily variation in the GnRH challenged T (T_GnRH_) levels, we conducted day and night sampling in each photoperiod. Day samples were taken 30 minutes after light onset and night samples were taken two hours after light offset (Fig 1 c, d).

### (e) Morphological Measurements, plasma collection and testosterone hormone Assays

We measured the cloacal protuberance volume (CPV), which enlarges in males due to sperm storage during breeding season and is used as a measure of gonadal growth leading to spermatogenesis. (Wolfson 1952). The volume of the CP was estimated using the equation for the volume of a cylinder, V= π(radius)^2^Height (Schut et al., 2012). The width (i.e., diameter) was measured at base of the CP using dial calipers and the height (H) was measured with a ruler. Postnuptial molt was scored at the end of the experiment based on primary feathers in both populations (Nolan et al., 2002). The progression of the 9 primary flight feathers molt at the end of breeding was calculated by allocating a score to each feather from 1-5 depending on the extent of molting feathers: an old unmolted feather 0 (0%), a primary missing or pin feather 1 (1-10%), a one-quarter grown feather 2 (11-30%), a one-half grown primary feather 3 (30-70%), a three-quarters grown primary 4 (70%-85%), and a completely new feather 5 (90%-100%). The scores derived from the percent feather growth were summed to generate a total molt score for each bird (Nolan et al., 1992; Ramenofsky et al., 2017).

We captured free-flying indoor birds to collect day and night blood samples. The birds received an intramuscular GnRH injection (~50 μl) into pectoral muscle of a dose of commercially available 62.5 μg / kilogram body mass of chicken GnRH (American peptide, Sunnyvale, CA) dissolved in PBS vehicle, which is known to activate the HPG axis in juncos (Jawor et al., 2006; Greives et al., 2016). Thirty minutes following the GnRH injection, a blood sample (50 μl) was taken from the wing vein to measure GnRH-challenged testosterone (T_GnRH_) levels. All birds were kept in an opaque bag between injections to reduce stress. Blood samples were immediately transferred to 4 °C fridge until later processing to extract plasma. Plasma samples were stored at – 20°C until processed for hormone extraction and quantification.

We determined GnRH-challenged testosterone (T) concentration from 20 μl plasma aliquots following established methods for our species (Jawor et al., 2006; Fudickar et al., 2016). Plasma samples were extracted three times using diethyl-ether prior to running hormone assays. After hormone samples were dried using nitrogen gas, each sample was dissolved in 250 μl of assay buffer. We used standard testosterone kit (Enzo Life Sciences, ADI-900-065, Farmingdale, NY) to determine circulating levels of T. All samples were measured in duplicate and randomized over twenty-eight plates. The inter-plate and intra-plate coefficient of variation were 7.32 ± 2.17% (mean ± SE) and 12.6% respectively.

#### Rate of change (k) in day T_GnRH_

We calculated rate of change in CPV day and night T_GnRH_ following Farner and Wilson (1957) and Follett and Maung (1978).

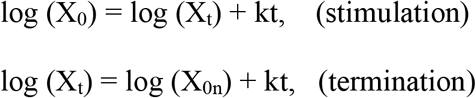

Where, X is reproductive response measure under increasing photoperiod (CPV, day and night T_GnRH_). X_0_ is the response at the beginning of the treatment, X_t_ is the peak stimulation response at t number of days, X_0n_ is the termination response at t number of days from the peak, and k is the logarithmic response growth rate constant/ day. The slope (k) gives a quantitative measure of the rate of change in gonad stimulation (initial point to peak) or termination (peak to endpoint) in a given light treatment calculated for a specific population.

### (f) Statistical Analysis

All data were analyzed using R (version 3.2.0). We used a Box-cox test of transformation to determine the normal distribution of all the response variables (i.e., CPV and day and night T_GnRH_ levels). We used a logarithmic transformation for CPV, day, and night T_GnRH_ levels. To test whether light treatment (F1), sub-population (F2), and photoperiod (F3), or interactions between the factors (F1, F2, F3) had a significant association with response variables, we used Multiple analysis of variance (MANOVA) followed by Tukey’s post-hoc multiple comparison tests (alpha < 0.05). In addition, we also used a generalized liner mixed-effect model (GLMM) for repeated measures for the same individuals, to ask whether light treatment, population, and day length as main effects or their interactions had significant impact on different physiological responses. To quantify whether light treatment and population, or the interaction between light treatment and population had a significant association with hydrogen isotope ratios values (δ^2^H) and molt score, we used two-way analysis of variance (2-way ANOVA) followed by Tukey’s post-hoc multiple comparison tests (alpha < 0.05). To assess co-variation between morphological measurements and δ^2^H isotope values as a continuous variable, we also performed linear regression for CPV, daytime T_GnRH_, molt score, rate of change (k) derived from CPV, day, and night T_GnRH_. We calculated the T_GnRH_ area under the curve (AUC) for each population and light treatment and calculated 95% confidence interval (CI). The non-overlapping CI were called significant of AUC comparison.

## 2. Results

### Hydrogen Isotope

Mean δ^2^H differed significantly between resident and migrant juncos (F_1, 32_ = 82.66, p < 0.0001; 2way ANOVA), but not between light treatments (Fig 1b). The δ^2^H (mean ± SE) was significantly lower in migrants than in residents (δ^2^H; migrants _NDL_ = −113.25%_0_ ± 4.31, migrants LAN = −107.33%_0_ ± 5.42, residents _NDL_ = −61.16%_0_ ± 9.91, residents _LAN_ = −50.67%_0_ ± 3.62_;_ Fig. 1b).

### Impact of ALAN on the timing of reproduction

CPV varied significantly by light treatment and photoperiod (F_12, 383_ = 15.4084, p < 0.0001; Fig. 2 a, b; Table 1). Molt score towards the end of the breeding showed significant effect of light treatment (F_1, 30_ = 59.94, p < 0.0001; 2-way ANOVA; Fig 2 c, d; Table 1 a). Day time T_GnRH_ showed significant interaction between light treatment and photoperiod (F_12, 383_ = 17.550, p < 0.0001; Fig. 2 e, f; Table 1 a). Night time T_GnRH_ showed significant effect of light treatment, photoperiod, and between two populations (F_12, 383_ = 2.4677, p = 0.004; Table 1 a).

**Table 1a.**
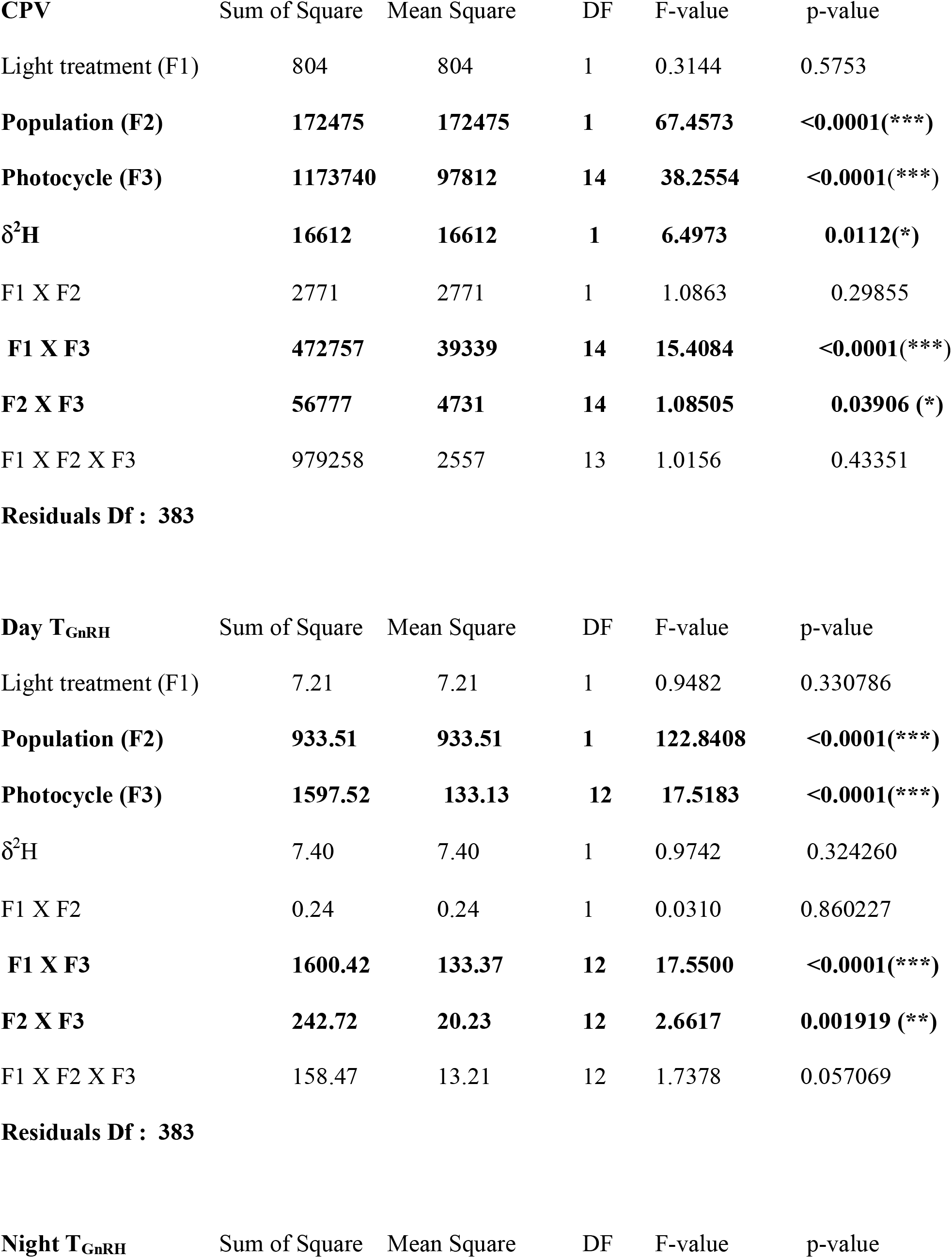
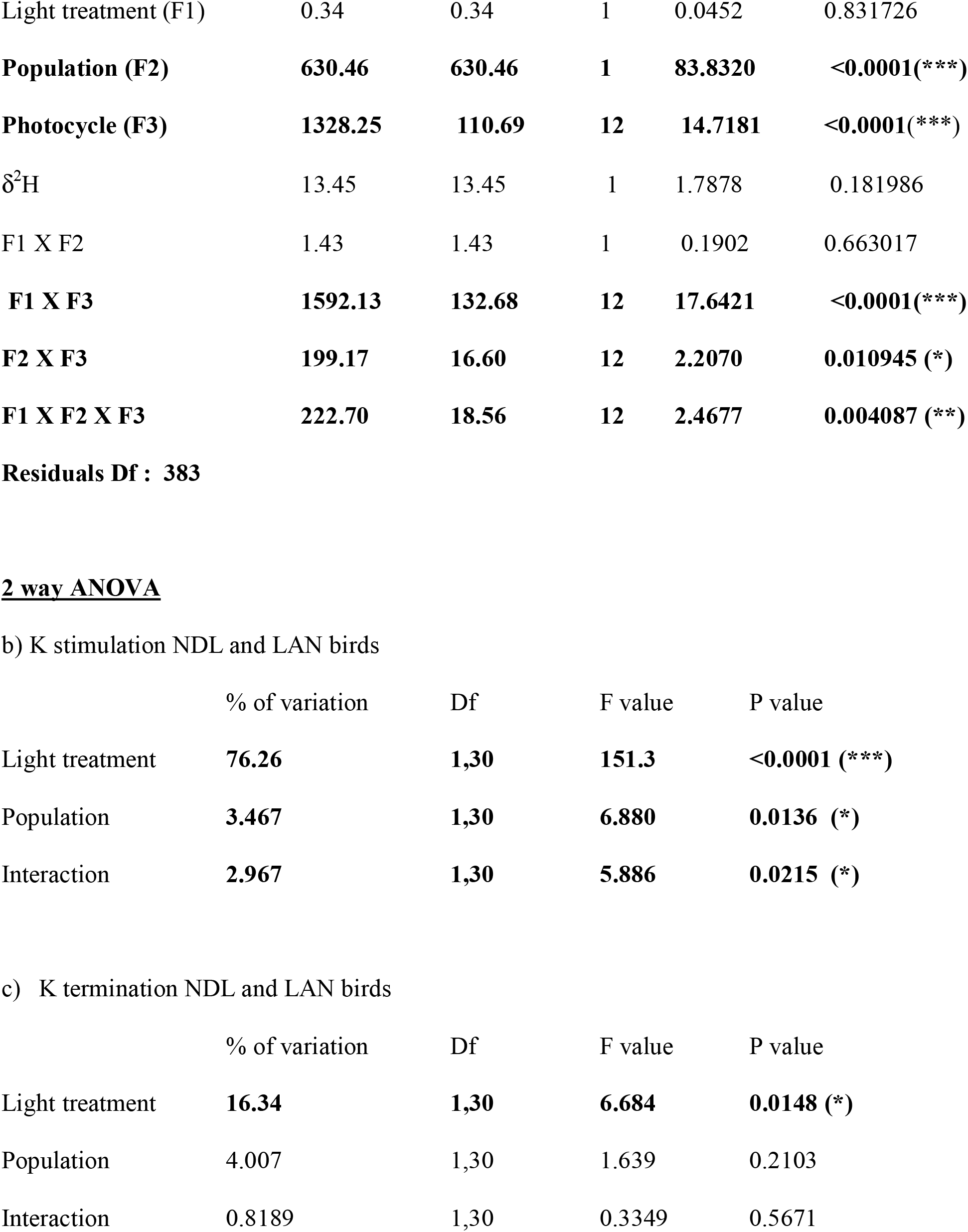
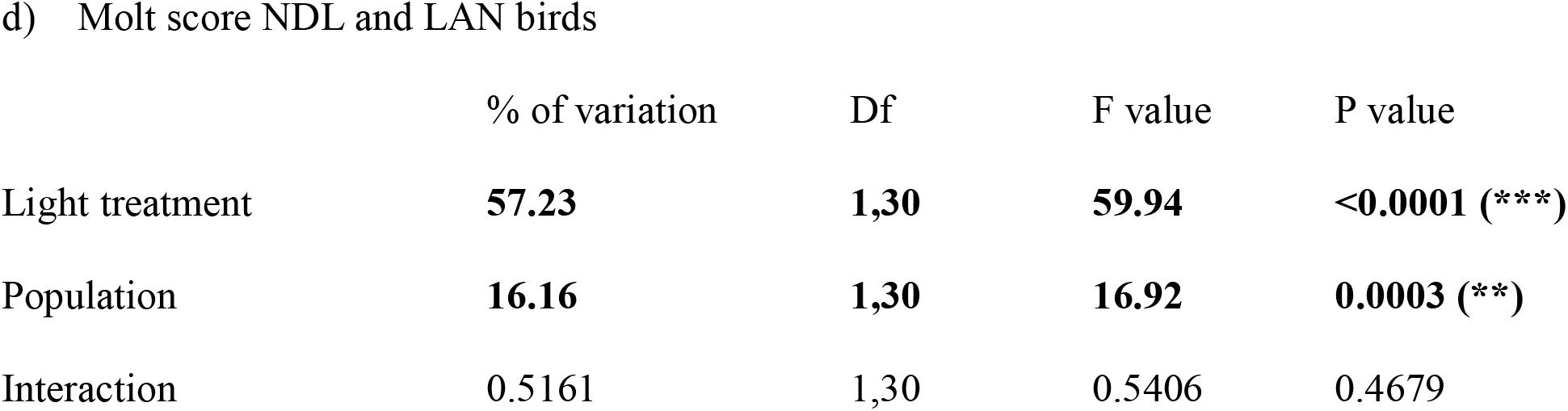
Factor (s) affecting different physiological responses (CPV, Day and night T_GnRH_): Multiple analysis of variance table followed by Tukey’s post-hoc multiple comparison tests (alpha < 0.05) derived from linear mixed effects model. Number of asterisk (*) denotes level of significance p <0.05(*), p <0.001(**), P<0.0001(***).

**Figure 2.**
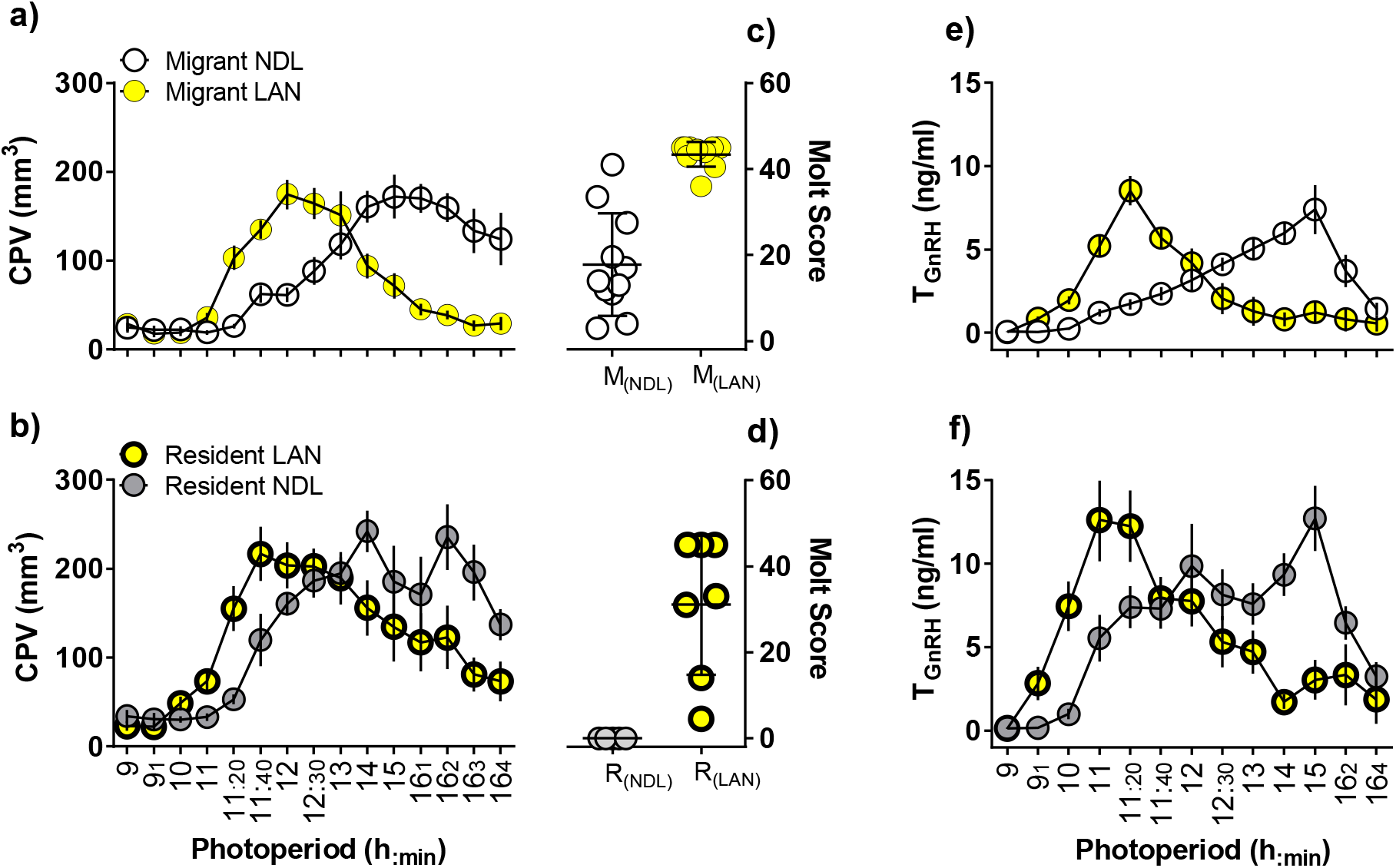
Impact of light at night treatment on cloacal protuberance volume, molt score, and GnRH challenged T (T_GnRH_). Measurements of CPV (a, b), molt score (c, d), and T_GnRH_ (e, f) in migrants (NDL: hollow circle; LAN: solid yellow circle) and residents (NDL: solid grey circle; LAN: thick solid yellow circle) starting from 9 h of light to 16h of light. Y-axis represents the CPV and T_GnRH_ response and X-axis represents increasing photoperiod in hours of day length exposure. Each data point represents mean ± SEM. Data were analyzed using a mixed-effect model with repeated measures for effects of light treatment, population, day length, and interaction between all three factors. Statistical significance was defined by alpha <0.05. Molt score was calculated only towards the end of the experiment.

Both resident and migrant birds elevated day time T_GnRH_ immediately after 2 weeks of ALAN exposure at 9 h of day length, whereas NDL birds did not exhibit increase in day time T_GnRH_ until 11 h of day length (Suppl. Fig. 1).

### Population specific impact of ALAN on the timing of reproduction

CPV growth showed significant effect of population and photoperiod (F_12, 383_ = 1.085, p = 0.039; Fig. 3 a, b; Table 1). The molt score showed significant difference in populations (F_1, 30_ = 16.92, p = 0.0003; 2-way ANOVA; Fig 2 c, d; Table 1 a). Irrespective of light treatment migrants showed heavy molt than resident population (p < 0.05; Tukey’s multiple correction). Day time T_GnRH_ showed significant interaction between population and photoperiod (F_12, 383_ = 2.6617, p = 0.001919; Fig. 3 e, f; Table 1 a). Similarly, night time T_GnRH_ also showed significant interaction between population and photoperiod (F_12, 383_ = 2.4677, p = 0.00408; Table 1 a).

**Figure 3.**
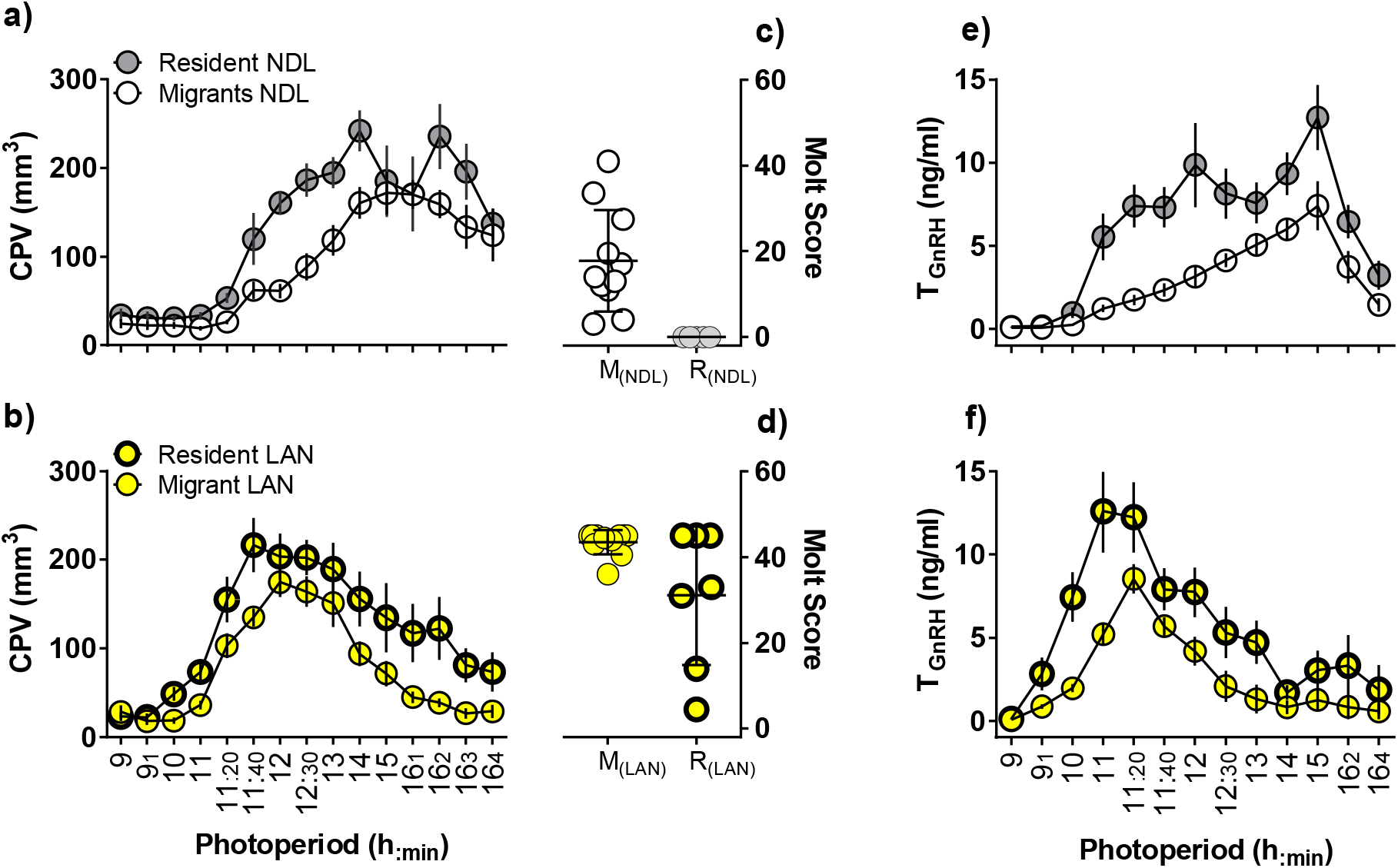
Population-specific response in cloacal protuberance volume, molt score, and GnRH challenged T (T_GnRH_) under different light treatments. Measurements of CPV (a, b), molt score (c, d), and T_GnRH_ (e, f) in NDL (migrant: hollow circle; resident: solid grey circle) and LAN (migrant: solid yellow circle; resident: thick solid yellow circle) starting from 9 h of light to 16h of light. Y-axis represents the CPV and T_GnRH_ response and X-axis represents increasing photoperiod in hours of day length exposure. Each data point represents mean ± SEM. Data were analyzed using a mixed-effect model with repeated measures for effects of light treatment, population, day length, and interaction between all three factors. Statistical significance was defined by alpha <0.05. Molt score was calculated only towards the end of the experiment. Data same as for Figure 2, plotted by population for easier comparison.

### Timing of reproductive initiation and termination to stable isotope values

We examined change in the relationships among CPV, Day T_GnRH_, with respect to δ^2^H values at 10L for initiation and at the end 16L for termination of breeding, considering residents and migrant collectively (Fig. 4)

**Figure 4.**
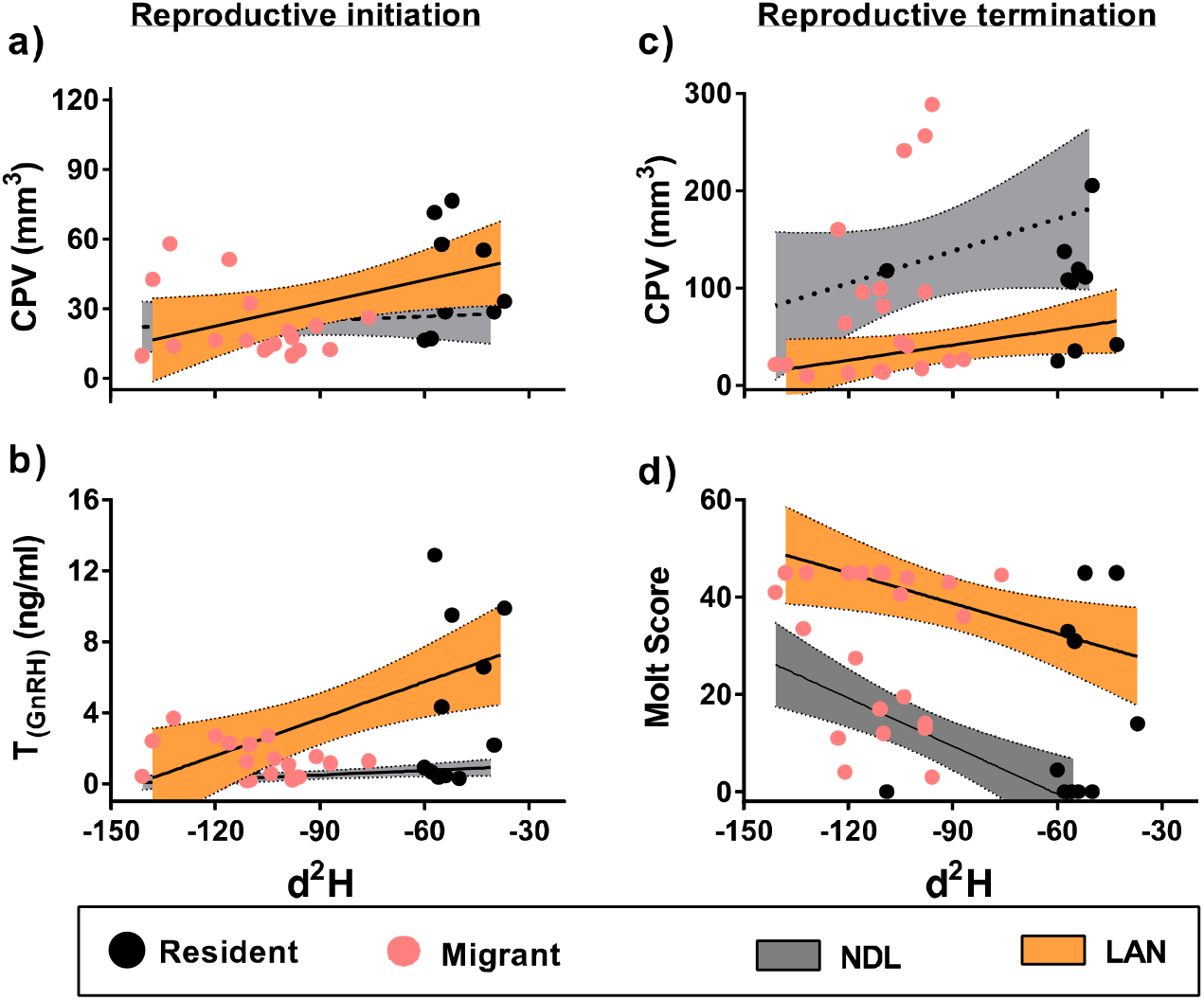
Latitudinal variation in timing of reproductive initiation and termination under NDL and LAN light treatments. Reproductive initiation measured in CPV and T_GnRH_ response in 10 L photoperiod and termination response in CPV and molt score during 16 L endpoint refractory state. Linear regression between NDL/ LAN CPV (a) and T_GnRH_ (b) response against δ^2^H as a constant variable under 10 L as reproductive initiation. Linear regression between NDL/ LAN in CPV (c) and molt score (d) response against δ^2^H as a constant variable under 16 L endpoint photoperiod as reproductive termination. X-axis represents the δ^2^H as a continuous variable and Y axis represents individual data points CPV, T_GnRH_, and molt score (migrant; solid pink circles. resident; solid black circle). Statistical significance was defined by alpha <0.05. Linear regression line with 95% confidence interval (shaded area) represents significant correlation as solid line and dotted line for no significant correlation.

**Figure 5.**
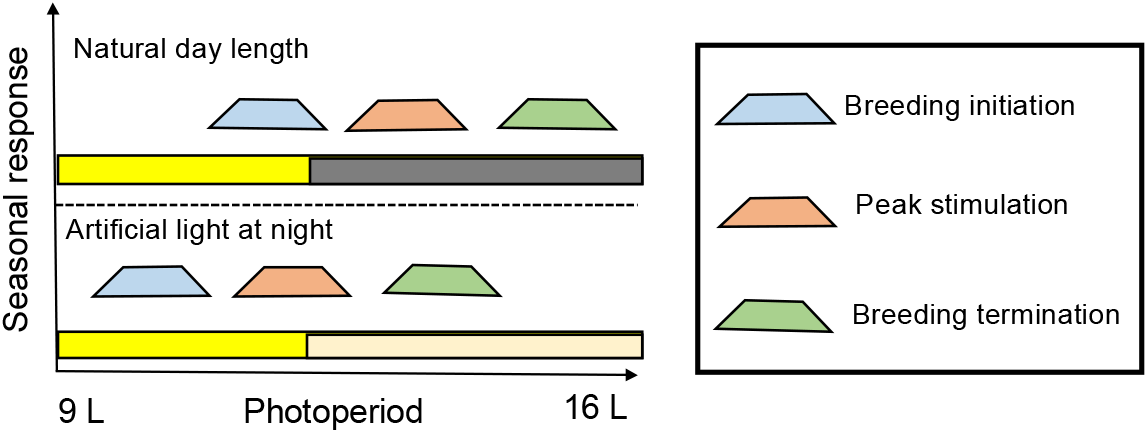
Schematic and simplified representation of effect of artificial light at night (ALAN) on different seasonal reproductive life-history states (LHSs). All LHSs: breeding initiation, peaks stimulation and termination are advanced in ALAN group.

#### Breeding initiation, and stable isotopes

During the initiation of breeding, δ^2^H values were significantly positively correlated with CPV (r = 0.4943, p = 0.037; Fig. 4a) growth in ALAN birds but not with NDL birds (r = 0.1436, p = 0.56; Fig. 4 a). Day time T_GnRH_ showed significant positive correlation in both the light treatment groups during recrudescence ALAN (r = 0.6102, p = 0.0072; Fig. 4 b) and NDL (r = 0.533, p = 0.027; Fig. 4 b).

#### Breeding termination, and stable isotopes

Birds under ALAN showed significant positive correlation with CPV (r = 0.567, p = 0.0175; Fig. 4 c) towards the end of the breeding. Whereas, birds exposed to NDL were still not regressed completely, hence δ^2^H values were not correlated with CPV (r = 0.388, p = 0.135; Fig. 4 c). When birds reached the refractory state molt scores (NDL; r = −0.742, p = 0.001, ALAN; r = −0.559, p = 0.0196; Fig. 4 d; Table 1 d) correlated with δ^2^H values providing evidence for latitudinal variation in timing of refractoriness.

##### Rate of change of reproductive stimulation and termination

We measured the rate of change (K) in gonad stimulation using day T_GnRH_ levels from start to peak level and termination from peak to end point. The K stimulation showed significant effect of light treatment and population (F_1, 30_ = 5.886, p = 0.0215; 2-way ANOVA; Table 1 b). K termination only showed significant effect of light treatment, day T_GnRH_ declined earlier in ALAN birds than NDL birds (F_1, 30_ = 6.684, p = 0.0148; 2-way ANOVA; Table 1 c).

##### T_GnRH_ Area under the curve (AUC)

Total AUC for T_GnRH_ varied significantly between resident and migrant birds irrespective of the NDL (resident _(mean±sem)_ = 77.16 ± 8.89, CI (59.72 – 94.6); migrant _(mean±sem)_ = 35.86 ± 5.19, CI (25.69 – 46.03) and ALAN (resident _(mean±sem)_ = 69.92 ± 9.87, CI (50.56 – 89.29); migrant (mean±sem) = 33.98 ± 5.39, CI (22.79 – 43.93) light treatments and the time of the day (Suppl. Table 1; Suppl. Fig 2 a-d)

## Discussion

The prodigious influence of increasing artificial light pollution is now a well-recognized potential threat to wildlife and biodiversity from almost all taxa (hölker et al., 2010; Gaston et al., 2013; Robert et al., 2015; Donners et al., 2018; de Jong et al., 2018; Dominioni et al., 2018). Using dark-eyed juncos as a model species, we investigated for the first time the effect of artificial light at night on the reproductive timing at population level, that is known to grow gonads at different critical threshold day length in a common garden experiment (Singh et al., 2019). Large effects on all the parameters that define the reproductive preparedness and termination were found advanced in birds exposed to ALAN. Migrant and resident birds exposed to ALAN induced their gonads, reached to peak stimulated phase earlier and terminated gonads sooner than their conspecifics exposed to dark nights. The breeding state measured as time between elevation and decline of T_GnRH_ levels remain unaffected between different light treatment in respective resident and migrant conspecifics. All the above observations indicate that a dim artificial light falling at night phase was perceived as a longer day length and accelerated reproductive response in both resident and migrant birds. In addition, we found that migrant birds maintained the phase lag in all the reproductive LHSs starting from gonad recrudescence, peak stimulation, and becoming photorefractory compared to resident population despite of different light treatments. This striking difference in the breeding timing at the population level irrespective of light treatment, suggests its regulation through a robust innate programs or differential regulation at the level of day length perception followed by induction of the neuroendocrine response.

### ALAN impact on reproductive timing

The annually changing light dark proportion over a 24 h day has the strongest effect in synchronizing the internal physiological response to the annual cycle (Helm et al., 2013). Studies suggest that ALAN could have profound impacts on desynchronizing seasonal reproductive timing to the external environment in wild inhabiting animals. The alignment of seasonal events such as annual migration, breeding, egg laying, and hatching are crucial to the food resources flux in wild. All these events are linked to each other and any misalignment in the one of the events might lead to the desynchrony in the linked seasonal states and annual cycle. Such misalignment in the reproductive timing and food resources has been shown to affect the migratory population predominantly, and can result in population decline at a large scale of around 90% in migratory pied flycatchers where nestlings provisioning time are misaligned with food resources flux (Both et al., 2006). The ecological consequences of breeding too early imposed reproductive costs on black-throated blue warblers (Lany et al., 2015). Nest survival probability may be lower early in the season if nests are easier to detect by predators when the leaves are not fully expanded, or if weather-related nest failures are more likely. A complementary explanation is that females which attempt to breed too early- before sufficient food is available to sustain that effort- pay a cost in body maintenance and capacity for subsequent nesting attempts (Nilsoon 1994, Perrins 1970).

In temperate region, species that breed north delay its breeding, have shorter reproductive seasons to synchronize nesting, egg laying, and nestling growth (Watts et al., 2015). Such populations with short breeding period corresponds to T elevation for shorter period (Singh et al., 2019). Hence, any perturbation in environment would affect the populations most that breed at higher latitude with short breeding period.

### Impact of ALAN on seasonal phenology at population level

Our study presents an insight into the impact of night light on the timing of different reproductive life-history states of two latitudinally distinct breeding populations of a north American sparrow the dark-eyed junco. We tested how exposure to night light will be perceived as long day length and accelerate the endocrine response which led to early breeding and terminate gonads sooner in a way that northern breeding population (i.e. migrants) is always later than the resident population breeding at lower latitude. This suggest that environmental cues at different breeding latitude could determine the breeding timing differences between the populations, which is maintained even if the reproductive response is accelerated by the external manipulation to the day length by introducing light at night. We observed early singing behavior along with spring migratory activity specific to migrants (based on observation, unpublished data) followed by early reproductive initiation in the birds exposed to ALAN, oppose to their counterparts experiencing dark nights in similar light dark schedule.

We suggest that annual seasonal phenologies are endogenously controlled by the mechanisms strictly adapted to maximize reproductive success. Thus, timing of breeding associated to latitude could be considered a key life-history event resetting the annual schedules in a long distance migratory bird. Field studies based on geolocator tracking data have demonstrated the influence of breeding latitude on migration schedules (Fraser et al., 2013; Stanely et al., 2015). Birds breeding in the temperate climate along the latitudinal gradient showed advanced phenology with the advancing spring (Lack 1950). Exposure to ALAN disrupts the day night cyclicity leading to desynchrony in the seasonal events. Dark eyed junco population breeding at higher latitude respond to longer CPP to activate HPG axis. Light falling to the dark phase accelerates the HPG response and advances the reproductive LHSs (Dominoni et al, 2013a; Dominoni 2015; Gaston et al., 2017). We cannot eliminate the possibility of difference in the molecular mechanisms at the level of deep brain photoreceptors, hypothalamic thyroid hormone metabolism, and consequent activation of HPG axis at population level that schedule the breeding time of populations across the latitude.

#### Reproductive response in ALAN comparable to Urban environment

ALAN treatment in our experiment provide a strong evidence how urbanization can advance timing of reproduction. The seasonal activation of HPG response is initiated by day length as predictive cue followed by supplementary cues (e.g., food, temperature, rainfall; Hahn et al., 2005; Wingfield et al., 2012; Watts et al., 2015). Advancement in breeding phenology might result from the plastic physiological response to adjust in novel urban environment. In a given scenario of long term occupancy of a population in urban environment might result birds to develop a separate niche with advanced breeding time might lead to reproductive desynchrony with birds occupying rural environment. This desynchrony in the reproductive timing as a result of difference in the sensitivity of HPG may limit gene flow between urban and rural populations. Alternatively, if the difference in the reproductive timing is a result of plastic adaptation to urban environment, then scope of divergence is limited. A study based on dark-eyed junco (*J. hyemalis thurberi*) population established recently (~35 years ago) in urban areas of San Diego CA USA, likely to diverge from an overwintering migratory population became sedentary with early breeding season (Atwell et al., 2012, 2014, Fudickar et al., 2017). A Study on wild population have shown that San Diego birds advance the first egg laying date by ~ 2.5 month earlier to the non-urban populations (Yeh and Price, 2004; Atwell et al., 2014). Further, when urban and rural junco populations housed in a common garden set up showed early elevation of baseline LH in urban juncos suggesting advanced endocrine response in urban birds (Fudickar et al., 2017). Similarly, urban population of Abert’s Towhees (*Melosone aberti*) in Phoenix Arizona USA, showed increased plasma LH than non-urban population (Davies et al., 2015). The early activation of urban birds endocrine response suggests that ALAN imposes persistent impact on the bird population that might restrict the gene flow between urban and rural birds and develop a separate niche. It has been observed that bird populations living in urban environment begin their reproductive development earlier compared to rural populations (Deviche and Davies 2013; Fudickar et al., 2017). Seasonal variation in the resource availability and climatic factors are often reduced in the urban environment, which might lead to a separate niche for birds formerly linked to migratory lineages that developed year-round occupancy in urban areas (Partecke and Gwinner, 2007; Atwell et al., 2014; Fudickar et al., 2017).

#### Study limitations

A limitation to our experimental design is that both resident and migrant birds experienced the similar rate of change of day length with maxima at 16 h of light in a common garden environment. Therefore, we cannot eliminate the possibility that migrants experience a more sudden increase in the day length while migrating to northern latitude with different peak day length ranging from 16 to 24 h of day light in late spring and early summer. The rate of gonadal growth (k) which is a measure of responsiveness to a given photoperiodic treatment in gonad growth may not be comparable for two populations in similar environment and therefore does not represent the ecologically relevant information. On the other hand, our common garden experiment allowed us to compare the difference in the overall reproductive response at population level under different light treatments. In our experiment the difference in the area under the curve for total duration of T_GnRH_ level varied significantly in two populations irrespective of light treatment, suggesting an inherent difference in the mechanisms regulating the timing of gonad stimulation, maintenance, termination and magnitude of peak gonad response specific to population. The more northerly population has longer critical photoperiod, delay breeding and terminate sooner than birds breeding at lower latitude (Dawson 2013; Fudickar et al., 2016; Singh et al., 2019). The difference in the timing of seasonal phonologies can be viewed as adaptations to schedule the reproductive timing of different geographic populations to keep the internal physiology in sync with the local environment most suitable to breed and hatchling provisioning.

We also recognize that NDL group simulating the natural change in photoperiod had dark nights may not necessarily allow for a direct comparison with natural photoperiod, where birds do experience celestial lights at night. It has been known that elevated corticosterone (CORT) levels have negative effect on reproductive system of birds (Wingfield and Sapolsky 2003). However, a captive study on peahens (*Pavo cristatus*) showed habituation to some extent under ALAN (Yorzinski et al., 2015). In our study, extended delayed in the peak stimulation of migrant juncos under NDL does suggest a possibility of HPG axis suppression contributing to extended delay in peak stimulation. However, although it remains to be studied in future how complete darkness / light at night in captivity affects the HPG axis response over a long period of exposure.

## Conclusion

Urban areas heavily equipped with ALAN sources on communication towers, street lights, skyscrapers, or lighthouses have been observed in attracting nocturnal migratory species across broad geographic regions (Falchi et al., 2016). The considerable amount of time spent in urban areas by migratory bird species during spring and autumn migration are more likely to get exposed to ALAN (La Sorte, Tingley, & Hurlbert, 2014). As the need to match with increasing food supply demand is increasing with population, majority of natural lands are increasingly converted into urban areas with ALAN sources to match up convenient human life style (Falchi et al., 2016; Gaston, Visser, &Hölker, 2015b; Kyba et al., 2017). The energy demanding migration and reproduction in birds is initiated by quest for food resources often funnels birds into human inhabited productive agricultural lands that also happen to be urbanized area with dense ALAN. This increasing encroachment by human habitat with ALAN and birds exposure to this novel environment has direct physiological impacts, such as altering the reproductive endocrine hormones, stress hormones, immune function, melatonin hormone levels, and sleep (Ouyang et al., 2015; 2017; 2018; Dominoni et al., 2013a; 2013b). The endocrine system that regulates the seasonal reproductive response is sensitive to the environmental changes and particularly to any external light source falling beyond the critical photoperiod phase in a 24 h daily light-dark cycle.

Combining information on both laboratory and field manipulations of ALAN will develop a better understanding for ecological consequences of the increasing urban environment on population diversity. We recommend that future studies investigate explicitly for the possibility of both phenotypic plasticity and evolutionary responses that may allow different populations to adapt well in a novel environment that ALAN creates.

## Ethics

All sampling methods were approved under protocol (# 15-026-17) by the Indiana University Institutional Animal Care and Use Committee.

## Authors’ contributions

D.S. and E.D.K. conceived the idea. D.S. and J.M. designed the study, collected samples. D.S. analyzed the data and produced the final draft with the help of J.M. and E.D.K.

## Acknowledgements

We thank Indiana Statistics Consulting Center, IU Bloomington for statistical help. We thank all Ketterson lab members for their suggestions in experiment designing and feedback on the manuscript. We thank David Sinkiewicz for providing access to the Center for Integrative Study of Animal Behavior (CISAB) lab facility.

## Supplementary figure and Table

**Figure S1.**
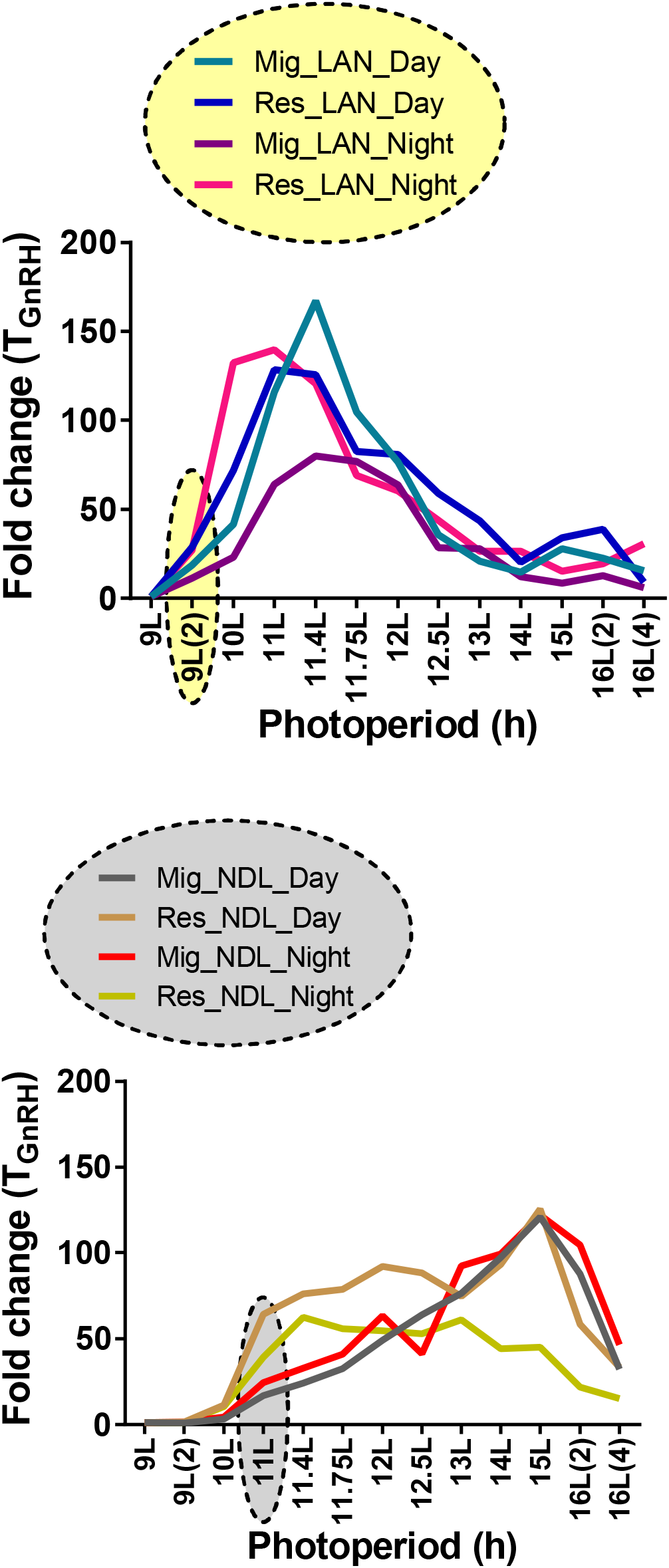
Line diagram to show the first day length at which mean fold change for day and night T_GnRH_ elevated.

**Figure S2.**
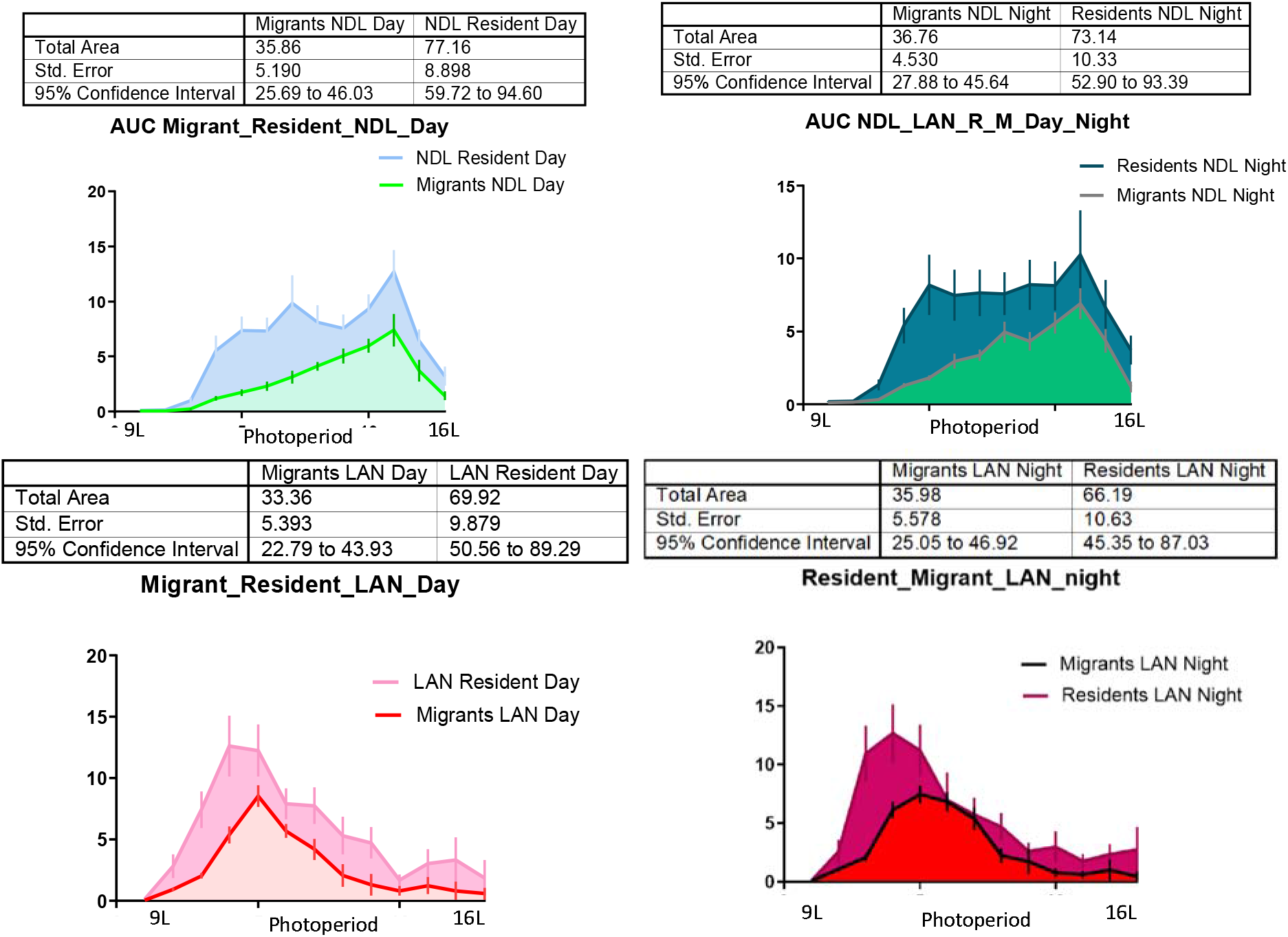
Area under the curve (AUC) calculated for day and night T_GnRH_ comparison between sub-populations and light treatments (a-d). The non-overlapping CI were called significant of AUC comparison.

